# THOC1 complexes with SIN3A to regulate R-loops and promote glioblastoma progression

**DOI:** 10.1101/2024.09.24.614748

**Authors:** Shreya Budhiraja, Shivani Baisiwala, Sia Cho, Rafal Chojak, Hasaan A Kazi, Amelia Stepniak, Ella N Perrault, Li Chen, Cheol H. Park, Crismita Dmello, Peiyu Lin, Adam M Sonabend, Atique U Ahmed

**Affiliations:** Department of Neurological Surgery, Feinberg School of Medicine, Northwestern University; Northwestern Medicine Malnati Brain Tumor Institute of the Lurie Comprehensive Cancer Center, Feinberg School of Medicine, Northwestern University; Northwestern University, Evanston, IL, 60208

**Keywords:** Glioblastoma, THOC1, R-loops, whole-genome CRISPR screen, progression

## Abstract

Glioblastoma (GBM), the most common and aggressive malignant brain tumor in adults, has a median survival of 21 months. To identify drivers of GBM proliferation, we conducted a CRISPR-knockout screen, which revealed THO Complex 1 (THOC1) as a key driver. Knocking down THOC1 significantly reduced GBM cell viability across patient-derived xenograft (PDX) lines, enhancing survival (p<0.01) in primary PDX models. Conversely, overexpressing THOC1 in non-cancerous cells bolstered viability, decreasing survival and causing tumor engraftment in vivo (p<0.01). Further investigation revealed THOC1’s robust interaction with SIN3A, a histone deacetylase complex. Histone deacetylation has been previously shown to prevent the buildup of R-loops, structures that form normally during transcription but can be lethal in excess. We found that THOC1-knockdown leads to elevated R-loop levels and reduced histone deacetylation levels. Next, to understand the networks specifically regulated by THOC1-mediated R-loop prevention, we conducted unbiased RNA-sequencing on control and THOC1-knockdown GBM cells. We found that THOC1’s role in R-loop prevention primarily affects telomeres, critical regions for cell replication. We further show that THOC1-knockdown results in significantly increased telomeric R-loop levels and shortened telomeres. Ultimately, this study suggests that targeting THOC1 shows promise as a therapeutic strategy to disrupt the delicate R-loop landscape and undermine GBM’s replicative potential.

## INTRODUCTION

One of the major hallmarks of cancer is its ability to facilitate great loads of cellular machinery required for uncontrolled cell division. One such machinery is R-loops, RNA: DNA hybrids that are formed normally during transcription. While R-loops are known for its usual role in regulating gene expression, unscheduled R-loop accumulation has been shown to pose a threat to genomic stability through the generation of DNA double-stranded breaks [2]. Given the genotoxicity of R-loop accumulation, the question of how cancer manages to maintain such high rates of replication in the presence of increased levels of R-loops without compromising its genomic integrity is poorly understood. Thus, elucidating how cancer cells avoid deadly R-loop accumulation is necessary for uncovering how this disease process is so aggressive.

Glioblastoma (GBM), in particular, is the most aggressive and common primary malignancy of the central nervous system in adults. Even with the multimodal standard of care treatment involving surgical resection, radiation, and temozolomide (TMZ)-based chemotherapy, GBM only has a median survival of approximately 21 months [1].

To better understand which genes promote and support GBM progression, our lab performed a CRISPR-Cas9 whole genome screen identifying key genetic drivers of this process. THO Complex 1 (THOC1), known to be part of the transcription-export (TREX) complex that promotes coupling transcription to mRNA export from the nucleus following cotranscriptional assembly on nascent RNA transcripts, emerged as key driver. THOC1 deficiency has been previously shown to cause defects in mRNA export. Furthermore, there is mounting evidence that THOC1 is elevated in several cancers, including prostate, ovarian, and colorectal. However, the specific role of THOC1 in GBM remains unknown [3] [4] [5].

Here, we show that THOC1 plays a key role in GBM aggression and tumorigenesis via tight regulation of R-loops. More specifically, the results of our study suggest that inhibiting THOC1 activity not only significantly increases genotoxic R-loop burden, but it does so to reduces tumor burden in vivo and prolong survival. The R-loop-mediated mechanism we propose here represents a new pathway for how glioblastomas maintain replicative potential and avoid DNA damage, something that can help us understand why the primary disease is so aggressive and why it is so resistant to classically cytotoxic therapies – cementing THOC1 as a promising therapeutic target in GBM.

## METHODS

### CRISPR-Cas9 knockout screen

The screening procedure closely followed previously established protocols [6]. We initiated the process by infecting H4 human GBM cells with the Brunello whole-genome knockout library (Addgene, Cambridge, MA, USA), which included around 19,000 genes, each with 4 sgRNAs, along with 10,000 sgRNA non-targeting controls. To prepare the library, we cultured 80% confluent HEK293T cells in a T225 flask for 20-24 hours. Subsequently, we introduced Opti-MEM I reduced serum, psPAX (10.4 µg/ml), pMD2.G (5.2 µg/ml), and Lipofectamine Plus reagent to the cells. After a 4-hour incubation, we filtered the media using 0.45 µM filters and stored the virus at -80 °C.

To determine the virus titer, we seeded 3 million H4 cells and 2 ml of media in a 12-well plate. We then added varying volumes of virus (ranging from 25 ul to 400 ul) along with 8 µg/µl of polybrene to each well, followed by spinfection at 1000 g for 2 hours at 33 °C. After a 24-hour incubation at 37 °C, we harvested the cells, seeded them at 4000 cells per well, and cultured them for 96 hours alongside non-transduced cells. Finally, we conducted a titer glo assay to assess cell viability and determine the multiplicity of infection (MOI).

Next, we utilized 70,000 sgRNAs to culture and spinfect 500 million H4 cells. Following a 4-day selection period with 0.6 µg/ml puromycin, 150 million cells remained viable. We extracted genomic DNA (gDNA) from 50 million of these cells after amplifying sgRNA with unique barcoded primers serving as controls. The remaining cells were expanded to 200 million over a 4-day period before undergoing treatment with 700 uM DMSO and TMZ for 14 days. Subsequently, we harvested the cells to amplify sgRNA and create the sequencing library.

The gDNA extraction was carried out using the Zymo Research Quick-DNA Midiprep Plus Kit (Cat no: D4075, Irvine, CA, USA), followed by purification with 100% ethanol containing 1/10 volume of 3 M sodium acetate (PH 5.2), and the addition of a 1:40 glycogen co-precipitant (Invitrogen Cat no: AM9515). To measure the gDNA concentration, we employed NanoDrop 2000 (Thermo Scientific, Waltham, MA, USA) and performed PCR for DNA amplification.

For sequencing, we used a NextSeq machine, generating 300 million reads for the four sgRNAs pool, each with 1000 reads/sgRNA. The sequencing process included 80 cycles of read 1 (forward) and 8 cycles of index 1, following the Illumina protocol. To enhance coverage, we introduced 20% PhiX into the NextSeq.

To analyze the computational data, we utilized CRISPRAnalyzR and the CaRpools pipeline. Significance of changes was determined using DESeq2 and the MaGeCK algorithms.

### Cell lines and culture

Patient-derived xenograft (PDX) glioma cell lines (GBM43, GBM6) were obtained from C.D. James at Northwestern University and maintained according to the published protocol [7]. In this case, the PDX cells were cultured in Dulbecco’s Modified Eagle’s Medium (DMEM) supplemented with 1% fetal bovine serum (FBS; Atlanta Biologicals, Lawrenceville, GA, USA) and 1% penicillin-streptomycin (Cellgro, Herdon, VA, USA; Mediatech, Herndon, VA, USA).

Human glioma cell lines, U251 and H4, were procured from the American Type Culture Collection (ATCC; Manassas, VA, USA). These cells were cultured in DMEM containing 10% FBS and 1% penicillin-streptomycin mixture. A frozen stock was used to replenish cells that had been expanded for a maximum of 4 passages. These frozen stock cells were stored at -180C in liquid nitrogen in pure FBS containing 10% DMSO (dimethyl sulfoxide).

Neural stem cell, astrocyte, and fibroblast lines were acquired from American Type Culture Collection (Manassas, VA, USA) and from Kerafast (Boston, MA, USA). H1B.F3 and fibroblast (WI-38) lines were cultured in DMEM supplemented with 10% FBS and 1% penicillin-streptomycin. NSC LM008 cells were cultured in neurobasal media (ThermoFisher Scientific; San Jose, CA, USA) and supplemented with B27 (no Vitamin A; Invitrogen, Carlsbad, CA, USA), N2 (Invitrogen), 1% penicillin-streptomycin, basic fibroblast growth factor (bFGF, 10ng/mL; Invitrogen), and epidermal growth factor (EGF, 10ng/mL; Invitrogen). To ensure optimal growth, media was replaced every 2 days.

### Animals and in vivo models

Athymic nude mice (nu/nu; Charles River, Skokie, IL, USA) were used in this study and housed according to Institutional Animal Care and Use Committee (IACUC) guidelines. All applicable federal and state statues governing the use of animals for biomedical research were also in compliance with this study.

Intracranial implantation of GBM cells was performed as previously described in our laboratory’s established GBM mouse model [8]. Animals initially received a prophylactic intraperitoneal (i.p.) injection of Buprenex and Metacam, followed by an anesthetizing i.p. injection of a ketamine/xylazine mixture (Henry Schein; New York, NY, USA). Sedation was confirmed by a toe-pinch test. Artificial tears were applied to their eyes, and the scalp was sterilized using Betadine and ethanol. Using a scalpel, the skull was exposed, at which point an approximately 1-mm burr hole was drilled above the right frontal lobe. The mice were then placed into a stereotactic rig where they received an injection of 150,000 GBM PDX cells using a Hamilton syringe 3mm from the dura, for a duration of 1 minute. To ensure proper release of the cell suspension, the needle was slightly raised and left undisturbed for 1 additional minute. Careful removal of the needle followed, and at this point the animal’s head remained in position while the skin of the scalp was sutured closed (Ethicon; Cincinnati, OH, USA).

Drug treatment regimens were initiated 7 days after intracranial implantation. I.p. injections of either TMZ (2.5mg/kg) or equimolar DMSO were delivered to each animal daily for 5 consecutive days. Following IACUC and Northwestern University guidelines, animals were sacrificed once it was evident that they would not survive past the following morning.

### Cellular transfection

To generate lentiviral particles, low-passage X293 cells (ATCC; Manassas, VA, USA) were plated at 80% confluency based on a previously described protocol [7]. 6 hours after plating, the cells were transfected with a mixture of HP DNA Transfection Reagent (Sigma-Aldrich; St. Louis, MO, USA) diluted in OptiMEM medium (Gibco; Waltham, MA, USA), in addition to packaging and target plasmids in accordance with the manufacturer’s instructions (Addgene).

### Viral transduction

48-72 hours after maintaining the cells following transfection, the virus supernatant was harvested and sterilized with a 45-µm filter and ultracentrifuged at 133,897 relative centrifugal force (RCF) for 2 hours to isolate viral particles. The resulting viral pellet was resuspended in phosphate-buffered saline (PBS) and aliquoted for storage in -80C until use. When ready for cellular transduction, cells were resuspended in 50µL of media, and 20 MOI lentivirus amounts were added per sample along with 1mL of appropriate media. This virus-media mixture was spun for 2 hours at 37C at 850g. Following centrifugation, the cells were plated with fresh media along with the viral media they were spun in. 48-72 hours later, the media was changed to non-viral containing media and the efficiency of the transduction was assayed via Western blot.

### Cell viability assays

Viability assays were performed according to a previously established protocol [9]. Cells were plated at 5000 per well in a 96-well plate with eight replicates per condition. Cells were allowed to grow and attach for 24 hours before treatments began. At this time, TMZ was administered using our laboratory’s standard dose-response protocol [7]. Following the assigned hours of treatment, the medium was removed and 110µL of MTT solution was instead added to the cells and the plate was then incubated at 37C for 5 hours. The MTT solution was prepared by diluting MTT stock reagent at 5 mg/mL in PBS. This mixture was diluted in fresh media at a ratio of 1:10 before adding it to each well. Post incubation, the medium was carefully removed, and the cells were resuspended in 100µL of DMSO to dissolve any crystals that had formed. After 10 minutes at room temperature, the place was loaded into a plate reader at an absorbance of 570nm and the data was analyzed to find percent viability in each well.

### Immunofluorescence

Mouse brains were harvest and frozen in cryoprotectant on dry ice, stored at -80C in 8µm sections, and stained according to standard immunohistochemistry protocol [7]. Sections were thawed at room temperature for 15-20 minutes and washed two times for 5 minutes in PBS + 0.05% Tween 20 (PBS-T) to remove residual cryoprotectant. The immunopen was then used to circle each brain section. Each section was fixed in ∼100uL of 4% PFA (ThermoFisher Scientific; Rockford, IL, USA) at room temperature for 15 minutes and washed twice for 5 minutes each time. Following this, antigen retrieval was performed by first boiling the samples in sodium citrate buffer for 20 minutes, then a 30-minute cooling period at room temperature. Slides were then washed 3 times for 5 minutes in PBS-T. Blocking was achieved by incubating slides in a 10% bovine serum albumin (BSA) solution with Triton-X (ThermoFisher Scientific; Rockford, IL, USA) for 2 hours at room temperature. Next, slides were incubated at 4C overnight with 100µL primary antibody solutions diluted in 1% BSA+ Triton-X. In the morning, the slides were washed 3 times for 10 minutes in PBS-T. 100µL of secondary antibodies diluted in 1% BSA + Triton-X were then applied and incubated for 2.5 hours at room temperature. Lastly, they were washed in PBS-T 3 times for 10 minutes. In order to image these slides, a drop of ProLong Gold Antifade reagent with 4’,6-di-amidino-2-phenylindole (DAPI) was added to each section (ThermoFisher Scientific; Rockford, IL, USA). Images of these slides were organized and assessed in ImageJ.

Additional experiments were conducted following the immunocytochemistry protocol previously published [10]. After removing chamber slides from incubation and washing once with PBS, 200µL of 4% PFA was added to each well and allowed to incubate at room temperature for 10 minutes before washing gently with PBS. Then, the cells were blocked for 2 hours at room temperature with 200µL 10% BSA solution. After aspirating the BSA off the slides, 100µL of primary antibody diluted in 1% BSA + Triton-X was added and the chamber was then incubated overnight in 4C. The next morning, the wells were washed 3 times for 5 minutes using 1% BSA + Triton-X. After this, 200µL of secondary antibody in 1% BSA + Triton-X was added and incubated at room temperature for 2 hours. Following incubation, the wells were washed 3 times for 10 minutes with PBS. Finally, ProLong Gold Antifade reagent with DAPI was added to each section to allow for imaging using a Leica microscope. These images were analyzed in ImageJ.

### Flow cytometry

Cells containing the plasmid of interest were harvested and spun down to a pellet, then washed with 100µL of PBS. Primary antibodies in FACS buffer (50µL per well) were added at room temperature in the dark and incubated for 1 hour. Cells were then spun down and washed once more with 100µL of PBS. Following this, the cells were resuspended in 80µL of FACS analysis buffer and spun down again. 100µL of fix perm buffer was added to each well. This buffer was prepared in a 1:3 ratio of fix/perm to buffer. Samples were then incubated at room temperature in the dark for 20 minutes. At this time, a 1:10 ratio of fix perm buffer was prepared in double-distilled water. 100µL of this solution was subsequently added on top of fix/perm. A 10-minute room temperature incubation period followed. Next, the samples were spun down for 5 minutes at 1500rpm and washed with the 1:10 perm buffer solution. The primary antibody was added to the perm buffer solution and 50µL was added to each well and allowed to incubate overnight at 4C. The next day, the samples were washed 3 times with FACS buffer and a secondary antibody contained in FACS buffer was added to them. The samples were incubated for 1 hour at room temperature. Finally, the samples were washed and resuspended in 100µL of FACS buffer and analyzed on the BD LSRFortessa Cell Analyzer.

### Western blotting

Cells were trypsinized, washed with PBS, and resuspended in mammalian protein extraction reagent (M-PER; ThermoFisher Scientific; Rockford, IL, USA) according to the protocol [11]. M-PER was supplemented with protease and phosphatase inhibitor cocktail (PPI’ ThermoFisher Scientific; Rockford, IL, USA) and EDTA (ThermoFisher Scientific; Rockford, IL, USA). Cells were vortexed for 1 minute 3 times with 10 minutes of rest on ice between each vortex. The resulting lysate solutions were centrifuged at 13,000rpm for 10 minutes at 4C. The supernatant was carefully collected, and the resultant protein concentration was determined via bicinchoninic acid assay (ThermoFisher Scientific; Rockford, IL, USA). This concentration was used to specify the amount of lysate required to make each Western blot sample. Each sample comprised of equal amounts of protein and varying amounts of SDS buffer (SDS sample buffer; Alfa Aesar; Ward Hill, MA, USA) with M-PER, resulting in equal total volumes across the samples. After a brief period of vortexing, the samples were boiled at 95C for 10 minutes before storing them in -20C until further use.

Samples were run through 8% SDS-polyacrylamide (SDS-PAGE, made in-house) via by gel electrophoresis (Bio-Rad; Hercules, CA, USA). The proteins were then transferred onto 0.45-μm polyvinylidene difluoride membranes (Milipore; Darmstadt, Germany). After transferring, the membranes were washed 3 times for 10 minutes in PBS then blocked for 2 hours with tris-buffered saline (TBS), comprised of 5% powered nonfat milk and 0.05% Tween 20 (Sigma-Aldrich; St. Louis, MO, USA). Next, the membranes were washed again 3 times for 10 minutes in TBS-T and subsequently placed in primary antibody solutions comprised of 5% BSA solution with sodium azide. The blots were then incubated overnight on a shaker at 4C. The next morning, the membranes were washed in TBS-T 3 times for 10 minutes, and placed in secondary antibody diluted 1:4000 in 5% milk. After an additional round of TBS-T washes, the membranes were coated in enhanced chemiluminescence (ECL, Clarity ECL, Bio-Rad). These membranes were developed using the ChemiDoc Imaging System (Bio-Rad).

### Dot blotting

To detect R-loops, cultured cells were trypsinized, pelleted, and washed with PBS to remove residual culture medium, followed by a previously validated protocol [12]. Briefly, the harvested cell suspension was transferred to 1.5 mL tubes. Subsequently, cold cell lysis buffer was added to the cell pellet at a rate of 300 μL per 2 × 10^6 cells, and thorough resuspension was achieved through pipetting. The cells were incubated on ice for 10 minutes and then centrifuged at 500 × g for 5 minutes to pellet the nuclei. The supernatant was discarded, and the nuclear pellet was resuspended in 400 μL of cold nuclear lysis buffer, followed by an additional 10-minute incubation on ice.

The subsequent purification of genomic DNA, which included RNA-DNA hybrids, commenced with the addition of 3 μL of 20 mg/mL proteinase K and an incubation period of 3–5 hours at 55 °C. The purification steps included DNA extraction using phenol:chloroform:isoamyl alcohol, precipitation, ethanol washing, and air drying. The resultant pellet was resuspended in 12 μL of elution buffer and further incubated for 30 minutes at 37 °C, and DNA concentration was subsequently measured via spectrophotometry. Nucleic acid samples were then diluted to desired concentrations in elution buffer. Positively charged nylon membranes were prepared, allowing 2 μL of each sample to be spotted. The samples were allowed to saturate into the membrane for at least 2 minutes before proceeding to UV crosslinking.

The membranes were incubated in blocking solution (5% milk in Tris-buffered saline with 0.05% Tween-20) for 1 hour at room temperature to minimize nonspecific binding. Subsequently, the membranes were incubated overnight in primary antibodies (in 5% milk in TBST) at 4 °C with shaking. Specifically, anti-dsDNA antibody (1:10,000 dilution) was added to one membrane, while the other received 1 μg/mL S9.6 antibody (1:1,000 dilution). Following incubation, the membranes underwent primary antibody removal and were washed three times with TBST for 5–10 minutes each. The membranes were then incubated with horseradish peroxidase (HRP)-conjugated secondary antibody (anti-mouse, 1:5,000 dilution) in 5% milk in TBST with shaking at room temperature. Subsequent to secondary antibody incubation, the membranes underwent three additional washes with TBST for 5–10 minutes each. Signal detection was achieved through ECL reagents, followed by signal intensity quantification using standard image processing tools such as ImageJ [12].

### Immunoprecipitation

Protein samples were isolated and normalized following the details above. A mixture of 5uL of ubiquitin antibody was incubated with 100µg of protein and protease/phosphatase inhibitors in Eppendorf tubes and placed in a rotary shaker in a cold room overnight. The next day, 30µL of Protein A/G beads were added to the sample and incubated for 2 hours at room temperature. Reactions were washed 3 times with 500µL of M-PER and centrifuged at 3200g for 5 minutes. After the third wash, 50µL of supernatant was reserved and 50µL of 2X SDS was added to all reactions. Samples were then incubated at 55C for 10 minutes to elute and were subsequently spun down again at 3200g for 5 minutes to separate the beads from the supernatant. Once the supernatant was collected, it was boiled for 10 minutes at 95C and loaded directly onto an SDS-PAGE gel. The remaining protocol was followed according to the Western blot protocol above.

### HDAC Activity Assay

The HDAC activity assay utilized an Abcam kit (ab156064, Cambridge, UK). Briefly, cells were gathered, and the lysate was obtained according to the manufacturer’s instructions. A black 96-well plate was prepared with the lysate and the provided kit buffers. Once the reaction commenced, the plate underwent continuous readings every 1-2 minutes using a plate reader.

### Telomere Isolation

Telomere isolation was utilized the TeloTAGGG™ kit (#12209136001, Millipore Sigma). Briefly, cells were gathered, and the genomic isolate was obtained according to the manufacturer’s instructions. Digestion of genomic DNA was then conducted using provided restriction enzymes, resulting in digestion of non-telomeric DNA and preservation of telomeric DNA and sub-telomeric DNA. Telomeric isolates were then utilized for further experiments.

### Quantitative PCR

Nucleic acid extraction was initially carried out according to previously discussed protocols. Following a 1:10 dilution of the isolated DNA in distilled water, reactions were arranged in triplicates, incorporating standardized quantities of DNA, SyberGreen (Thermo Fisher, Rockford, IL, USA), as well as forward and reverse primers (IDT, Newark, NJ, USA), which were intended for subsequent quantitative PCR analyses. The outcomes were recorded using a conventional qPCR apparatus. All primer sequences were generated using Primer-BLAST.

### Statistical analysis

Statistical analyses were performed and represented using GraphPad Prism v9.0 software (GraphPad Software; San Diego, CA, USA). Generally, data are presented as mean with standard deviation for continuous variables and number or percentage for categorical variables. Differences between two groups were assessed using Student’s *t* test or Wilcoxon rank sum test where applicable. Differences between multiple groups were assessed using ANOVA with *post hoc* Tukey’s test, or Mann-Whitney *U* test followed by Bonferroni correction as appropriate. Survival curves were graphed with the Kaplan-Meier method and compared by log-rank test. All tests were two-sided and p<0.05 was considered statistically significant. *In vitro* experiments were performed in biological triplicates.

## RESULTS

### Whole-genome CRISPR Screen Reveals THOC1 as a Major Driver of GBM Aggression

A whole-genome CRISPR-Cas9 knockout screen in human H4 GBM cells was performed to identify novel genes critical for driving GBM progression. The screen utilized a Brunello whole-genome knockout library covering over 19,000 genes and 200+ non-targeting controls using four sgRNAs per gene. Guides were sequenced at days 0, 14, and 28, and those discovered to be depleted – corresponding to genetic drivers of GBM – were further analyzed to identify top hits for genes responsible for the oncogenic progression (**Figure 1A**). Sequencing quality assessments confirmed the validity of screen results through appropriate per base sequence quality, per sequence quality scores, and sequence length distribution **(Figure S1A)**. Gene read count distributions and sgRNA frequencies showed expected differences across conditions, further confirming the reliability of the screen in identifying essential drivers of GBM through analysis of depleted guides **(Figure S1B)**. Finally, principal component analysis (PCA) assessment of similarities across replicates and differences across conditions substantiated screen quality **(Figure S1C)**.

**Figure 1:**
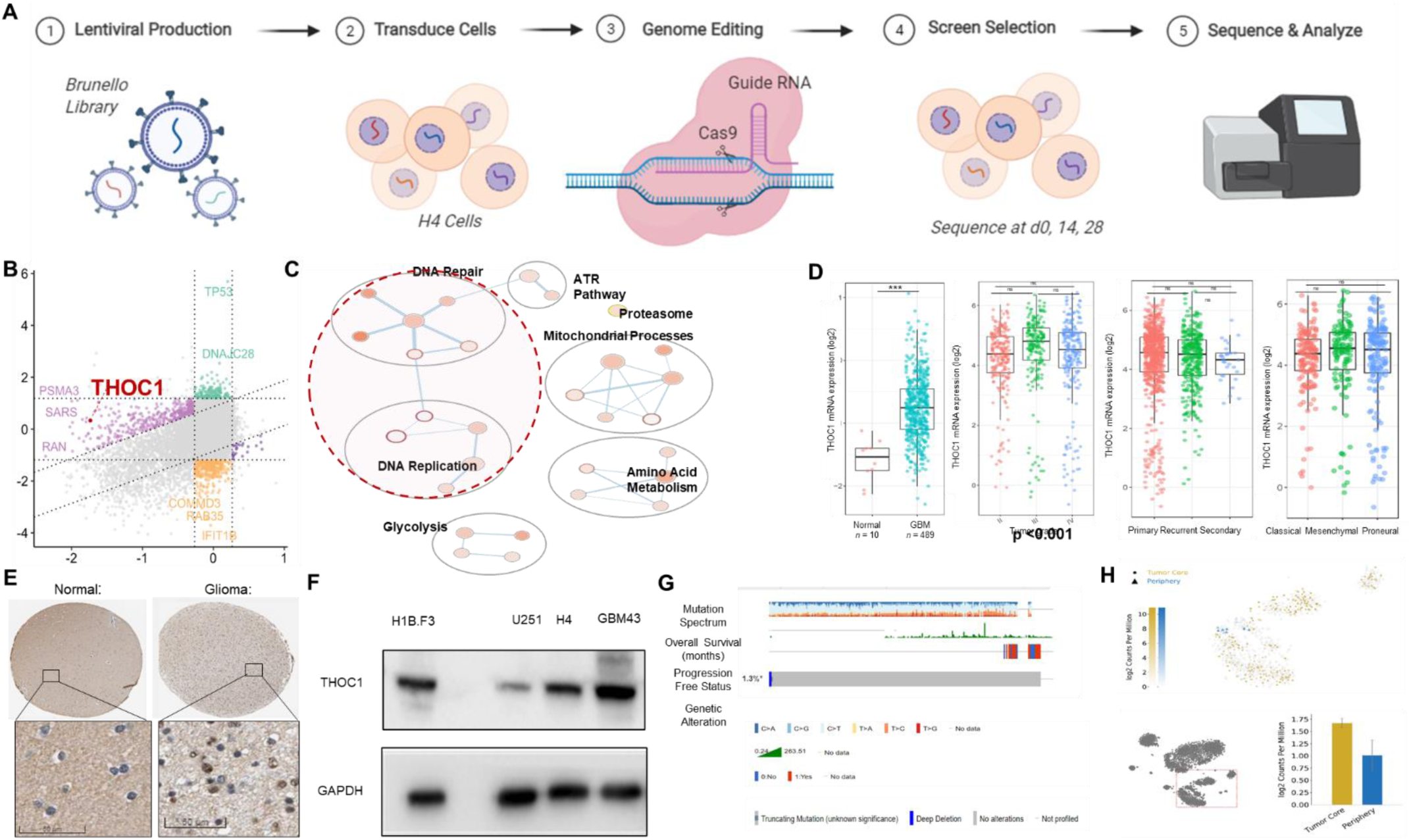
Whole-genome CRISPR Screen Reveals THOC1 as a Major Driver of GBM Aggression. ***A)*** Schematic of the whole-genome CRISPR-Cas9 knockout screen in human H4 GBM cells using the Brunello library to identify genes critical for GBM progression. Sequencing at days 0, 14, and 28 highlights top depleted guides as potential oncogenic drivers. ***B)*** Identification of THOC1 as a significant hit from the CRISPR screen. ***C)*** Pathway enrichment analysis showing THOC1’s involvement in several key pathways identified in the screen. ***D)*** Elevated THOC1 RNA expression in GBM compared to non-tumor samples, based on data from the GlioVis portal. ***E)*** Increased THOC1 protein expression in GBM tissue compared to normal brain tissue, as shown by Protein Atlas data. ***F)*** Western blot analysis showing higher baseline THOC1 expression in GBM patient-derived xenograft (PDX) line (GBM43) compared to a non-cancerous line. ***G)*** cBioPortal data indicating a low mutation rate of THOC1 in GBM (approximately 5%). ***H)*** Single-cell RNA sequencing data from GBMSeq showing high THOC1 expression localized to the tumor core.

THOC1 was of particular interest owing to its significant log fold change, novelty, and involvement in several pathways enriched in our screen **(Figure 1B, 1C, S1C)**. To begin, we looked at publicly accessible GBM patient datasets to better understand THOC1 expression in GBM. THOC1 RNA expression – obtained from GlioVis portal patient data – was observed to be elevated in GBM compared to non-tumor (p<0.001) across multiple datasets [14] (**Figure 1D**, **S2A, S2B)**. Similarly, Protein Atlas revealed increased protein expression in GBM tissue compared to normal brain tissue, as qualitatively assessed from patient samples [13] (**Figure 1E**, **S2C)**. However, there was no significant correlation between THOC1 RNA expression and tumor grade, recurrence status, or tumor subtype (**Figure 1E**, **S2A, S2B)**. This trend was also noted via western blot analysis, which revealed elevated baseline THOC1 expression in various GBM cell lines when compared to a non-cancerous line (**Figure 1F**). Slight differences in expression among GBM lines were also noted; thus, we proceeded to conduct all future experiments in multiple cell lines to account for subtype-specific characteristics (**Figure 1F**). Patient data from cBioPortal further revealed that THOC1 mutation rates in GBM are very low at around 5% [15] [16] (**Figure 1G**), and single-cell RNA sequencing data from GBMSeq showed that THOC1 expression highly localized to the tumor core [17] (**Figure 1H**). Moreover, THOC1 expression is higher in contrast-enhancing regions compared to non-enhancing areas **(Figure S2D)**.

It is important to note that THOC1 expression did not exhibit dependence on TMZ. GlioVis portal patient data revealed that patients who received standard-of-care temozolomide (TMZ) did not concurrently exhibit higher THOC1 RNA expression **(Figure S3A)**. Furthermore, single-cell sequencing analysis performed in our lab compared post-DMSO, post-TMZ, and mid-therapy GBM cells, revealing consistent RNA expression levels, indicating that THOC1 expression is not altered during therapy **(Figure S3B, S3C)**. These results were verified at the protein level, with western blot analysis showing no significant differences across varying time points after TMZ exposure, suggesting that THOC1 may be necessary for GBM progression but not specific to TMZ therapy **(Figure S3D)**.

### THOC1 Shows Effects on GBM Viability In Vitro and In Vivo

Patient survival curves from publicly available GBM datasets in GlioVis portal revealed that higher levels of THOC1 RNA expression were strongly correlated to shorter survival time for patients, corroborating its function in promoting GBM aggression and confirming the clinical relevance of our target as a potential therapeutic target (**Figure 2A**). To further validate this target, we then looked at the relationships between THOC1 and the well-known proliferation markers PCNA, Ki67, MCM2, and PLK1. Results indicated that these markers have substantial positive correlations with THOC1 RNA expression (**Figure 2B**).

**Figure 2:**
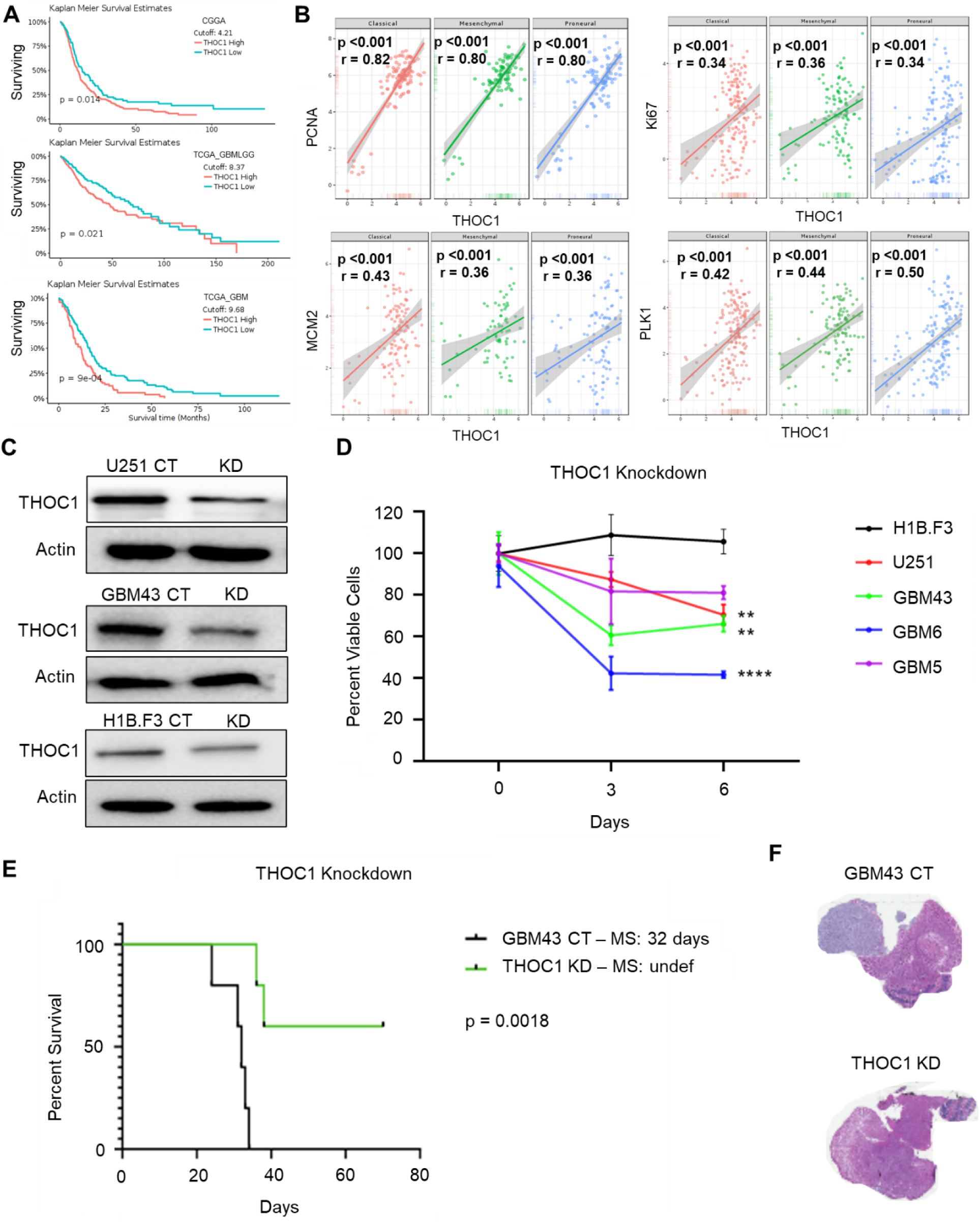
THOC1 expression is inversely correlated with survival. ***A)*** Kaplan-Meier survival curves from GlioVis portal showing that higher THOC1 RNA expression correlates with shorter survival in GBM patients. ***B)*** Positive correlations between THOC1 RNA expression and proliferation markers (PCNA, Ki67, MCM2, PLK1). ***C)*** Western blot confirming effective THOC1 knockdown in GBM cell lines using CRISPR-Cas9, with substantial reduction in THOC1 protein levels. ***D)*** MTT assays demonstrating significant decreases in viability in THOC1-knockdown GBM cell lines, while neural stem cells (H1B.F3) remain unaffected. ***E)*** Kaplan-Meier survival analysis of mice implanted with THOC1-knockdown GBM43 cells, showing significantly extended median survival compared to controls. ***F)*** H&E staining of mouse brain tissue showing a marked reduction in tumor size in mice implanted with THOC1-knockdown GBM43 cells.

Next, to investigate the role of THOC1 on GBM cells’ viability, THOC1-knockdowns were established using CRISPR-Cas9 constructs in H1B.F3, U251, GBM43, GBM5, and GBM6 cell lines. MTT assays were then conducted in normal and THOC1-knockdown lines to determine the effects on viability. As expected, three of the four GBM cell lines – GBM43, GBM6, and U251 – demonstrated significant decreases in viability (p<0.01) (**Figure 2D**, **S4A)**. In contrast, neural stem cells (H1B.F3) did not show a similar decrease in viability. The results of this cell viability assay were similarly verified with other sgRNAs, showing significantly reduced proliferation in THOC1-knockdown U251 and GBM43 cell lines when compared to THOC1-knockdown fibroblasts (p<0.05) **(Figure S4B).**

Next, mice were implanted with a GBM43 PDX THOC1-knockdown line and observed until they reached their endpoint. THOC1-knockdown resulted in a significant increase in median survival when compared to the control. While control mice had a median life of around 32 days, THOC1-knockdown mice had a significantly longer median survival (p<0.01) (**Figure 2E**). Following the death of the mice, brains were harvested and immunohistochemically stained for THOC1 to confirm knockdown **(Figure S4C)**. H&E staining of tumor tissue was also conducted to visualize tumor morphology. Indeed, tumor size was reduced dramatically in mice implanted with THOC1-knockdown GBM43 cells (**Figure 2F**).

### THOC1 Overexpression Results in Tumorigenesis

We then sought to determine whether, apart from driving growth of GBM, THOC1 may also transform normal neural stem cells when overexpressed to exhibit an oncogenic phenotype. ORF clones were used to overexpress THOC1 in three non-cancerous cell lines: H1B.F3 (a neural stem cell line), NSC LM008 (a neural stem cell line), and WI-38 (a fibroblast line). Western blotting confirmed the efficacy of THOC1 overexpression (**Figure 3A**). Preliminary experiments aiming to compare cellular architecture between control and THOC1-overexpression cell lines revealed a distinct morphological presentation of unique process-like features in THOC1-overexpression cells, perhaps suggesting a more migratory character associated with this cancerous phenotype (**Figure 3B**).

**Figure 3:**
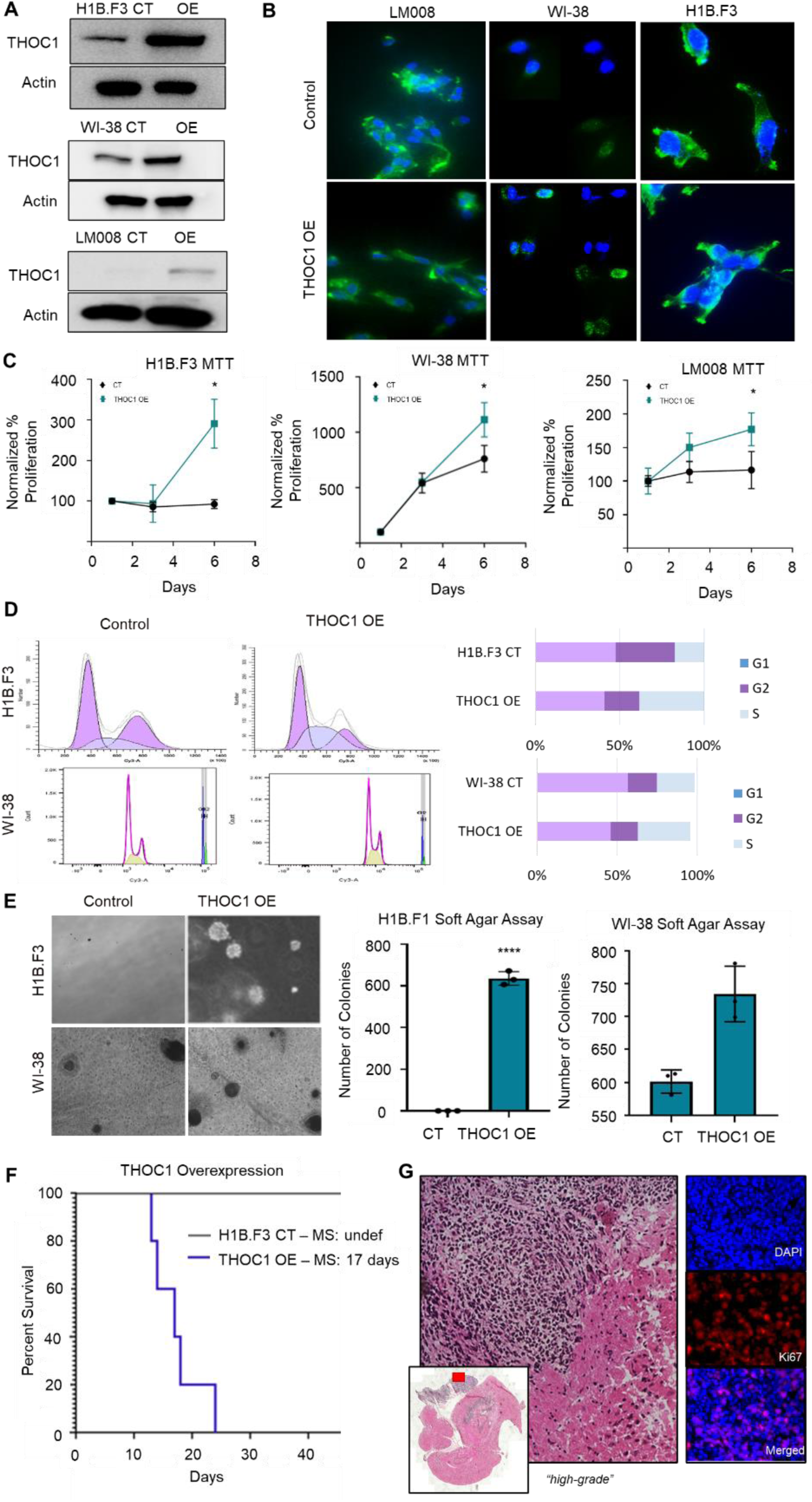
THOC1 expression promotes proliferation and migration. ***A)*** Western blot confirming successful THOC1 overexpression in three non-cancerous cell lines (H1B.F3, NSC LM008, WI-38). ***B)*** Morphological analysis showing distinct process-like features in THOC1-overexpression cells, suggesting a more migratory, cancerous phenotype. ***C)*** MTT assays demonstrating significantly increased proliferation in THOC1-overexpression cells across all three cell lines, with H1B.F3 cells showing the greatest increase. ***D)*** Cell cycle flow cytometry analysis showing a higher percentage of THOC1-overexpression H1B.F3 and WI-38 cells in the S-phase. ***E)*** Soft agar assay results showing large colony formation in THOC1-overexpression cells, particularly in H1B.F3, indicating transformation and anchorage-independent growth. ***F)*** Kaplan-Meier survival analysis of mice injected with THOC1-overexpression H1B.F3 cells, showing accelerated decline and shorter survival compared to controls. ***G)*** Histopathological analysis of THOC1-overexpression tumors in mice, showing high-grade tumor development and significant Ki67 staining.

Tumor proliferation of THOC1-overexpression cells was then measured via MTT assays and cell cycle flow cytometry. THOC1-overexpression cells in all three cell lines – H1B.F1, LM008, and WI-38 --demonstrated significantly higher cell proliferation than control cells (p<0.05) (**Figure 3C**). In particular, H1B.F3 cells showed the greatest difference, proliferating nearly 300 percent relative to the control, which did not experience as much growth within the 6-day timeframe (**Figure 3C**). Additionally, cell cycle flow cytometry analysis showed a higher percentage of cells in the S-phase for both H1B.F1 and WI-38 cell lines. THOC1-overexpression H1B.F3 cells contained 38.39% of its population in this phase, as compared to 17.07% in control H1B.F3 cells, while THOC1-overexpression WI-38 cells similarly showed 32.80% of its cells in this phase, as compared to 23.40% in its control cells (**Figure 3D**).

To examine the tumorigenicity in THOC1-overexpression cells, soft agar assays were performed. Soft agar assays have been classically used to determine if cells demonstrate transformative potential through the ability to grow in an anchorage-independent manner that is associated with cancer cells but not normal cells. Assays were conducted in both H1B.F3 and WI-38 lines, revealing that THOC1-overexpression cells demonstrated large colony formation. In H1B.F3 cells specifically, THOC1-overexpression cells formed around 600 colonies, as compared to none in the control (p<0.001) (**Figure 3E**).

We next wanted to assess whether THOC1-overexpression cells might develop tumors in vivo. H1B.F3 cells with control or overexpression lentiviral vectors were injected into mice, and animals were monitored for tumor development and end point survival. The brains of the mice were collected and sectioned for histopathology once they had achieved their endpoint. THOC1-overexpression cells resulted in accelerated decline and mortality when compared to controls, with THOC1-overexpression mice living an average of 17 days (**Figure 3F**). Histological examination of harvested THOC1-overexpression tumors by an in-house licensed neuropathologist revealed tumor development, which was classified as “high grade.” Tumors also exhibited significant Ki67 staining, reflecting their significant proliferative nature (**Figure 3G**). Overall, these findings robustly indicate that THOC1 overexpression demonstrates a capacity to induce a tumorigenic phenotype in vitro and in vivo.

### THOC1 Complexes with SIN3A to Prevent R-loop Formation in GBM

THOC1, or THO complex 1, canonically functions in the TREX (transcription/export) complex in order to regulate and ensure efficient export of polyadenylated RNA. However, recent literature has demonstrated a novel interaction between this gene and SIN3A, a multi-protein complex responsible for recruiting major HDACs 1 and 2 to facilitate transcriptional repression via local histone deacetylation [18].

It has been widely shown that deacetylation is a prime mechanism for the prevention of R-loops, structures that form normally during transcription [19]. Specifically, by transiently closing the chromatin after passage of the RNA polymerase, HDAC recruitment renders the DNA inaccessible and unsuitable for forming R-loops.

R-loops are ubiquitous three-stranded nucleic acid structures—consisting of a DNA:RNA hybrid that has displaced a ssDNA. Although these structures are notable for their typical function of regulating gene expression, R-loops have also been observed to have another critical function when unregulated and accumulated: to threaten genomic stability, generate DNA breaks, and cause cell death.

Given the ability of these structures to cause significant genomic damage, maintaining proper R-loop levels is paramount to cellular homeostasis. In fact, defects in R-loop equilibrium have been linked to several malignancies, where R-loop disequilibrium may drive oncogenic progression. Given the importance of R-loops in cancer and our identification of THOC1 as a promising therapeutic target, we postulated that THOC1 upregulation in GBM enables increased SIN3A-mediated deacetylation required for preventing excess R-loop formation and thus cell death (**Figure 4A**).

**Figure 4:**
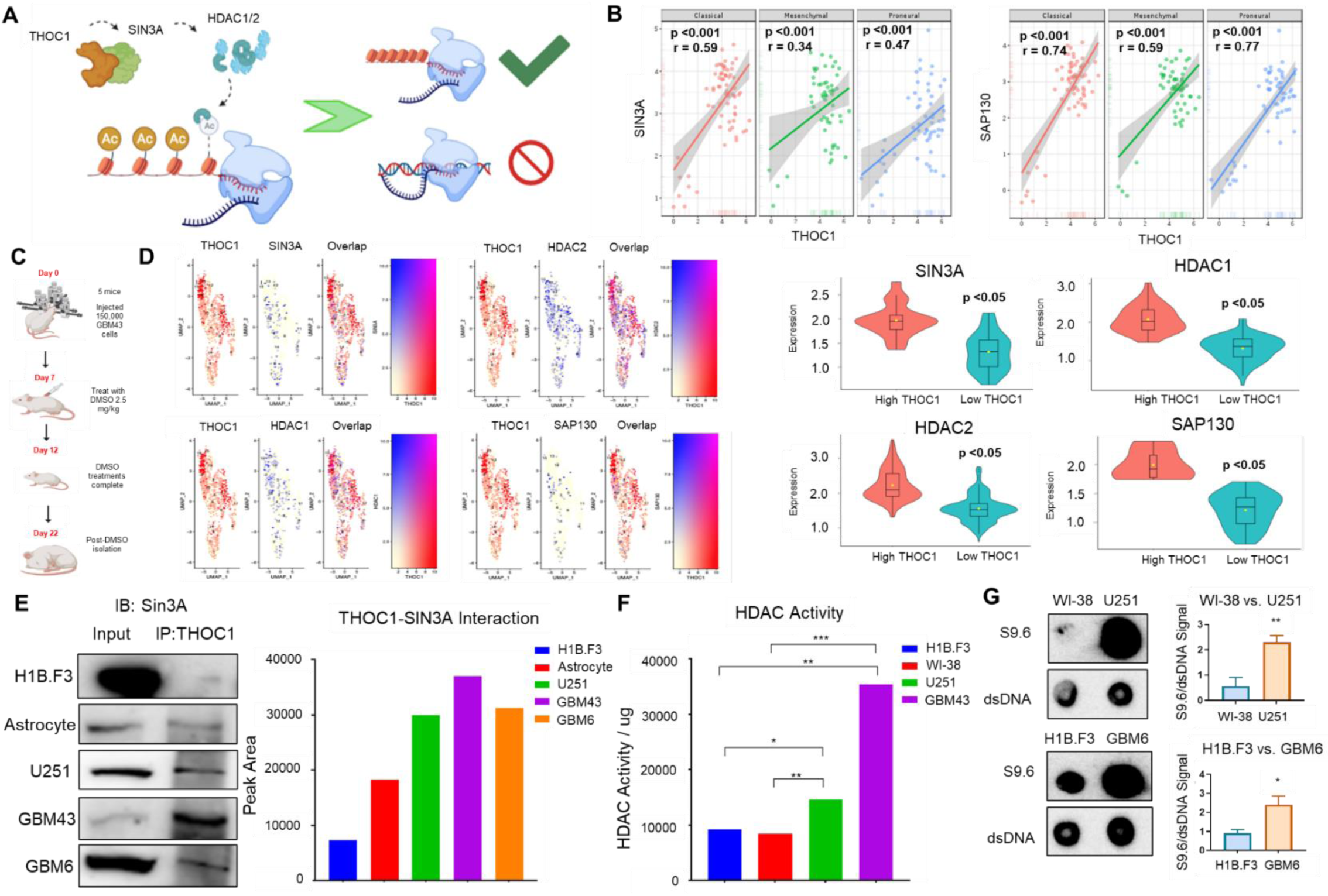
THOC1 Upregulation in GBM Enhances SIN3A-Mediated Deacetylation, Reducing R-loops and Promoting Cell Survival. ***A)*** Schematic representation illustrating the proposed mechanism by which THOC1 upregulation in GBM may lead to increased SIN3A-mediated deacetylation, preventing R-loop accumulation and subsequent cell death. ***B)*** Correlation analysis of patient data from the GlioVis portal showing strong positive correlations between THOC1 and SIN3A, as well as SAP130 mRNA expression levels (p<0.001). ***C)*** Single-cell sequencing of GBM43 cells implanted in vivo. ***D)*** Stratification analysis from single-cell sequencing showing the correlation of THOC1 expression with SIN3A and associated genes at a single-cell level. ***E)*** Immunoprecipitation of THOC1 in GBM and non-cancerous cell lines, showing greater THOC1-SIN3A interaction in GBM lines compared to neural stem cells and astrocytes. ***F)*** HDAC activity assays demonstrating higher HDAC activity in GBM cells compared to neural stem cells and fibroblasts (p<0.05). ***G)*** Dot blot analysis using the S9.6 antibody to detect DNA hybrids, showing greater R-loop levels in U251 and GBM6 cells compared to fibroblasts and H1B.3 neural stem cells.

To investigate this, we first wanted to determine whether THOC1 expression is correlated with SIN3A expression. Patient data obtained from the GlioVis portal revealed robust mRNA expression correlation of THOC1 and SIN3A, as well as SAP130, a known subunit of SIN3A (p<0.001) (**Figure 4B**). To further corroborate this correlation, we also performed single-cell sequencing, where GBM43 cells were implanted and then isolated and sequenced to see in vivo correlations (**Figure 4C**). THOC1 RNA expression was found to be highly correlated with SIN3A, SAP130, HDAC1, and HDAC2, as represented on a single cell level through stratification of low and high expression from optimal cutoff analyses (p<0.05) (**Figure 4D**). These strong correlations between THOC1 and genes associated with SIN3A thus amplify the importance of THOC1’s high baseline expression in GBM, supporting the potential significance of this altered histone deacetylation activity in promoting GBM aggressiveness.

We then wanted to confirm whether the previously reported interaction between THOC1 and SIN3A was present in GBM. Immunoprecipitation of THOC1 in a variety of non-cancer and GBM lines revealed greater THOC1-SIN3A interaction in all three GBM lines when compared to neural stem cells and astrocytes, supporting the notion that THOC1 upregulation in GBM may enable greater interaction with SIN3A and thus increased deacetylation levels (**Figure 4E**). In fact, to assess the effects of this increased interaction in GBM on HDAC activity, HDAC activity assays were performed, in which exogenous fluorophore-tagged peptides with acetyl groups fluoresced upon endogenous HDAC-mediated deacetylation [23]. Indeed, GBM cells were found to exhibit greater HDAC activity when compared to neural stem cells and fibroblasts (p<0.05) (**Figure 4F**).

After confirming that the elevated THOC1-SIN3A axis present in GBM allows for heightened levels of deacetylation, we then sought to determine whether this phenomenon is a mechanism to cope with the imminent threat of increased R-loop levels in GBM. To do so, we first attempted to confirm whether GBM displayed greater R-loop levels like in other cancers with similar transcriptional demand needed to sustain uncontrolled cell division. Dot blot analysis of nucleic acid extracts was conducted in various cell lines using the S9.6 antibody, a highly specific antibody for DNA:RNA hybrids. Greater S9.6 signal was seen in U251 and GBM6 cells when compared to fibroblasts and H1B.3 neural stem cells, respectively, supporting the notion that GBM may require THOC1-mediated deacetylation to avoid toxic R-loop accumulation (p<0.05) (**Figure 4G**).

### THOC1 Mediates Genomic Stability through Modulation of R-loops

Non-cancerous cells may not exhibit elevated THOC1-expression due to their low basal replication and R-loop levels. On the other hand, GBM cells may have increased THOC1-expression in order to combat the risk of accumulated, unscheduled R-loops that can potentially lead to cell death. Given this, we postulated that THOC1’s crucial role is to control this continual seesaw of transcriptional R-loops (allowing for deadly GBM proliferation) and closing of the DNA to avoid R-loop dysregulation (preventing DNA damage and GBM death) (**Figure 5A**).

**Figure 5:**
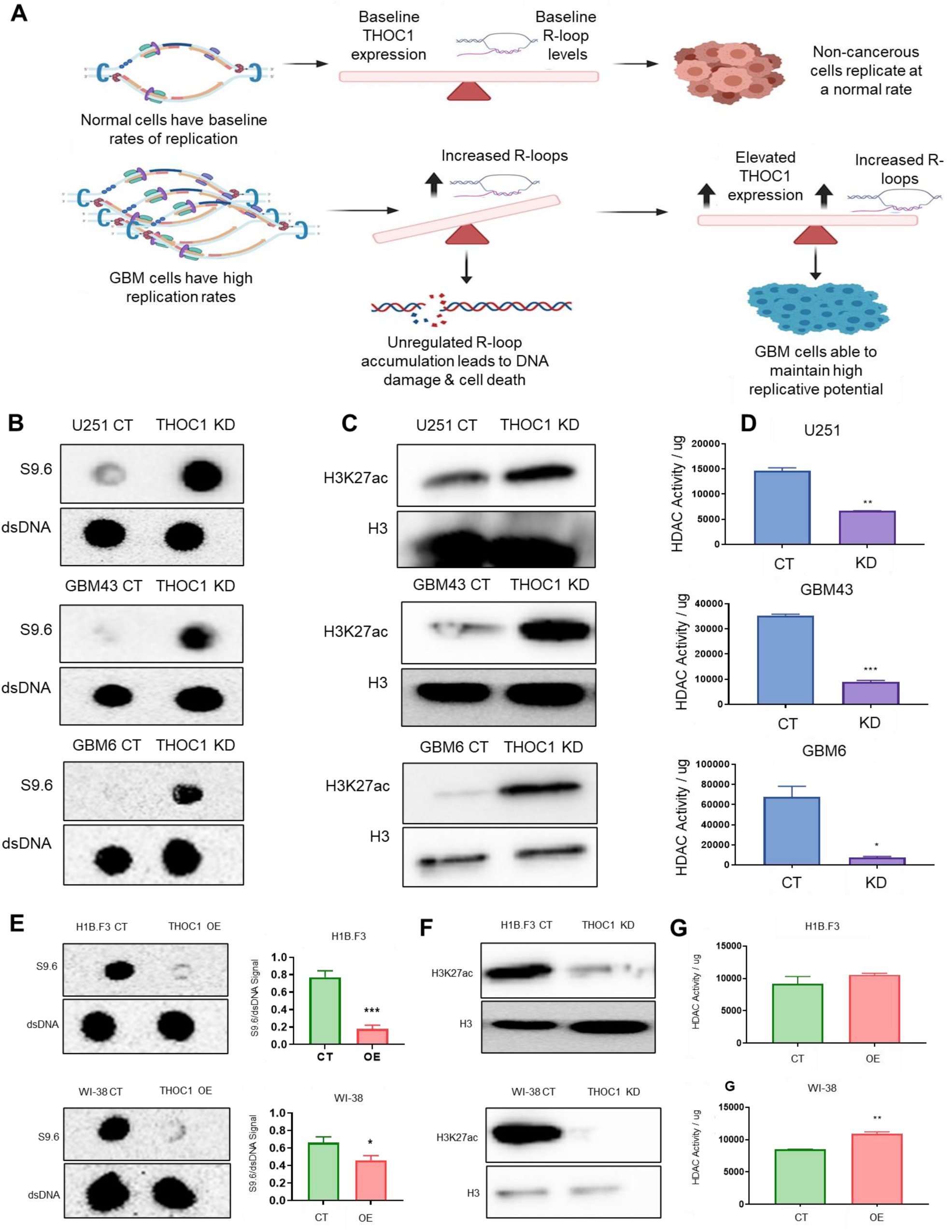
THOC1 promotes R-loops formation through modulation of HDAC activity. ***A)*** Schematic representation of the hypothesized role of THOC1 in controlling R-loop levels in GBM cells to balance transcriptional activity and prevent DNA damage. ***B)*** Dot blot analysis showing a significant increase in R-loop levels in THOC1-knockdown GBM cells (U251, GBM43, GBM6) compared to control cells (p<0.01). ***C)*** Western blot analysis indicating increased global acetylation levels (H3K27ac) in THOC1-knockdown GBM cells. ***D)*** HDAC activity assays demonstrating significantly lower deacetylation activity in THOC1-knockdown GBM cells compared to control cells. ***E)*** Dot blot analysis showing that THOC1 overexpression in non-cancerous neural stem cells and fibroblasts significantly decreases R-loop levels. ***F)*** Western blot analysis showing lower acetylation levels (H3K27ac) in THOC1-overexpressing non-cancerous cell lines. ***G)*** HDAC activity assays showing increased HDAC activity in THOC1-overexpressing non-cancerous cell lines, with significant increases observed in WI-38 fibroblasts (p<0.05) and similar trends in H1B.F3 neural stem cells.

We have previously shown that THOC1-knockdown causes a decrease in viability in GBM cells. Given our hypothesis, we believed this decrease in viability is due to the inability of THOC1 to handle the burden of high R-loop levels, leading to R-loop-associated DNA damage.

To investigate THOC1’s influence on R-loops, we first measured S9.6 in control and THOC1-knockdown GBM cells. In all three cell lines – U251, GBM43, and GBM6 – THOC1-knockdown resulted in a significant increase in R-loop levels, as compared to the control (p<0.01) (**Figure 5B**). To identify whether this increased R-loop burden was due to THOC1’s association with SIN3A, we next assessed whether THOC1-knockdowns resulted in global changes in acetylation levels via western blot analysis. Indeed, greater levels of acetylation, as observed by H3K27ac, were observed in the knockdown conditions for all three cell lines – U251, GBM43, and GBM6 – confirming THOC1’s unique role in GBM’s epigenetic landscape to influence R-loop formation (**Figure 5C**). HDAC activity assays similarly reflected the same phenomenon, with THOC1-knockdown GBM cells exhibiting significantly lower deacetylation activity (p<0.05) (**Figure 5D**).

Given that THOC1 also demonstrated the ability to promote tumorigenesis, we then sought to determine whether this THOC1-overexpression in non-cancerous lines would result in R-loop biology similar to GBM cells. Dot bot analysis revealed that THOC1-overexpression significantly decreased R-loop levels in both normal neural stem cells and fibroblasts (p<0.05) (**Figure 5E**). Furthermore, both lower acetylation levels and higher HDAC activity were concordantly observed in the THOC1-overexpression lines, as indicated by H3k27ac expression from western blot analysis and HDAC activity assays (**Figure 5F**, **5G)**. Of the two lines, THOC1-expression in WI-38 fibroblasts demonstrated significantly increased HDAC activity (p<0.05), while THOC1-overexpression in H1B.F3 neural stem cells showed a similar trend (**Figure 5G**).

### THOC1 enhances telomere stability through prevention of telomeric R-loops

Given that GBM cell lines may be overexpressing THOC1 to maintain R-loop homeostasis under increased proliferative stress, we hypothesized that THOC1-knockdown in GBM cell lines promotes histone acetylation, leaving GBM cells vulnerable to R-loop-mediated DNA damage and cell death. To better understand the epigenetic landscape of GBM cells with respect to R-loops, we performed bulk RNA-sequencing in control and THOC1-knockdown cells, followed by GSEA to find pathways particularly affected by decreased THOC1 activity to stabilize the R-loop landscape.

Analysis of pathways most downregulated in THOC1-knockdown cells revealed telomeric packaging, protein folding, and RNA polymerase I promoter opening as the most enriched pathways (**Figure 6A**). To further understand how THOC1 may influence telomere homeostasis via R-loops, we compared telomeric R-loop abundance in control and THOC1-knockdown cells.

**Figure 6:**
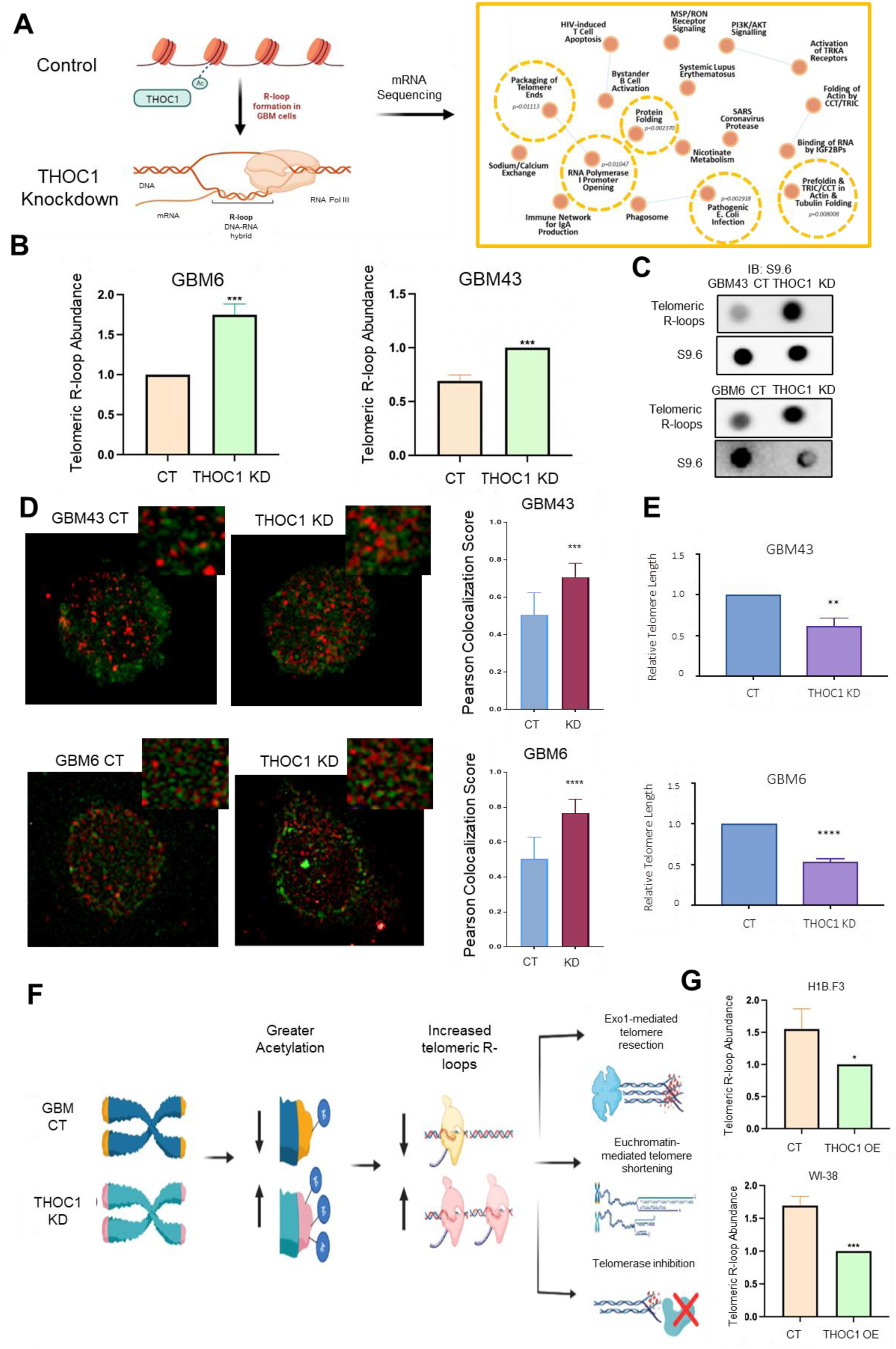
THOC1 enhances telomere stability through prevention of telomeric R-loops. ***A)*** GSEA analysis of bulk RNA-sequencing data from THOC1-knockdown and control GBM cells, revealing downregulated pathways including telomeric packaging, protein folding, and RNA polymerase I promoter opening in knockdown cells. ***B)*** Comparative analysis showing a significant increase in telomeric R-loop abundance in THOC1-knockdown GBM cells. ***C)*** Dot blot and immunoprecipitation analysis with S9.6 antibody, showing increased telomeric R-loops in THOC1-knockdown cells compared to control cells. ***D)*** Immunofluorescence staining of telomere marker (TRF1) and R-loops (S9.6) showing greater co-localization of TRF1 and S9.6 in THOC1-knockdown GBM cells compared to control cells, indicating elevated telomeric R-loop presence. ***E)*** Analysis of telomere length in THOC1-knockdown cells showing significantly reduced telomere lengths. ***F)*** Schematic model proposing that THOC1’s influence on GBM viability and progression may be predominantly driven by its role in suppressing telomeric R-loops rather than global R-loop levels. ***G)*** Analysis showing increased telomere length in THOC1-overexpressing neural stem cells and fibroblasts.

We observed a significant increase in telomeric abundance via qP in THOC1-knockdown GBM cells, confirming that THOC1-mediated R-loop prevention affects telomeres specifically (**Figure 6B**). We further validated these results via dot blot analysis of telomeric isolates, followed by immunoprecipitation of S9.6, in the same cell lines, which exhibited an increase in telomeric R-loops in THOC1-knockdown cells relative to the controls (**Figure 6C**). This was also seen via immunofluorescence staining of telomere marker (TRF1) and R-loops (S9.6), which showed significantly greater co-localization of TRF1 and S9.6 in THOC1-knockdown cells compared to control cells (**Figure 6D**).

Given this increased presence of telomeric R-loops in THOC1-knockdown cells and their implications on telomere damage, we next investigated whether increased abundance of these structures resulted in reduced telomere length, a proxy for overall telomeric health. Indeed, we observed that THOC1-knockdown cells exhibited significantly reduced telomere lengths, suggesting that THOC1 may play an even more crucial role in regulating the global R-loop landscape: specifically suppressing R-loops on telomeres to ensure telomeric maintenance (**Figure 6E**).

We have previously shown that THOC1’s influence on R-loops in GBM cells is the result of its role in the epigenetic process of deacetylation, which may be leading to increased telomeric R-loops as well. However, given the unique implications of R-loops specifically on telomeres – including the promotion of Exo1-mediated resection, euchromatin-mediated telomere shortening, and telomerase inhibition – perhaps the effect of THOC1 on GBM viability and progression that we see is predominately driven by its role in influencing telomeric R-loops specifically, rather than global R-loop levels (**Figure 6F**). Additional studies revealed that THOC1-overexpression in neural stem cells and fibroblasts results significantly reduced telomeric R-loop abundance and increased telomeric length in both cell lines, highlighting the possibility that similar mechanisms may be at play in THOC1’s role in tumorigenesis (**Figure 6G**, **Figure S5A).**

## DISCUSSION

One of the major hallmarks of GBM, like many other cancers, is its ability to thrive in the presence of genomic instability. This genomic instability is the driving force behind the cascade of stepwise, accumulated mutations that lead to its heterogeneous and aggressive phenotype. Within this context, R-loop-induced genomic instability emerges as a significant player, contributing to the intricate landscape of GBM progression and oncogenesis. However, the current role of R-loops in GBM is currently unknown, owing to its paradoxical nature. On one hand, they are instrumental in facilitating the rapid transcriptional demands required for uncontrolled cell division, which is a hallmark of cancer progression. In fact, R-loops serve as essential intermediaries in gene expression, playing a role in regulating various cellular processes. However, the paradox lies in the fact that, when in excess, dysregulated R-loop formation can pose a significant threat to genomic stability, resulting in cell death. Given that R-loops serve as a double-edged sword in cancer biology, our study aimed to uncover how GBM straddles this fine line between R-loop paucity and excess to promote GBM progression.

We first show that THOC1 is elevated in GBM compared to non-cancerous cell lines. Knocking down THOC1 resulted in significantly decreased GBM viability when compared to non-cancerous cell lines, suggesting its unique role in GBM progression. In vivo studies using a GBM43 PDX model with THOC1-knockdown cells revealed a significant increase in median survival for mice implanted with THOC1-knockdown cells compared to the control group. Notably, tumor size was dramatically reduced in mice implanted with THOC1-knockdown GBM43 cells, emphasizing THOC1’s role in tumor growth and aggressiveness. Apart from GBM progression, we then show the effects of THOC1 overexpression in non-cancerous lines. In vitro studies reveal that THOC1-overexpression resulted in increased cell proliferation, cell cycle progression, and anchorage-independent growth, highlighting THOC1’s ability to promote a tumorigenic phenotype. Moreover, in vivo experiments showed that THOC1-overexpressing non-cancerous cells was able to induce tumor engraftment in mice, demonstrating the oncogenic potential of THOC1 and its role in transforming normal cells into a cancerous phenotype.

To delve deeper into the functional aspects of THOC1 in GBM progression, we investigated its interaction with SIN3A, a histone deacetylase complex known to recruit HDACs 1/2, responsible for the majority of histone deacetylation activity. Histone deacetylation plays a pivotal role in preventing excessive R-loop accumulation by rendering closed DNA inaccessible to transcriptional machinery. We postulated that THOC1 may hinder R-loop formation by interacting with SIN3A, thereby stabilizing the R-loop landscape, preventing detrimental R-loop buildup, and enabling GBM proliferation.

We found that GBM cells display significantly greater levels of R-loops compared to non-cancerous cells, aligning with the idea that GBM cells have greater transcriptional demand (and thus require more machinery such as R-loops) to sustain its aggressive proliferation. Interestingly, we also found robust THOC1-SIN3A interaction in GBM cells compared to non-cancerous cells, which was similarly reflected by greater histone deacetylation activity in GBM. These results suggest that the elevated THOC1 expression in GBM may enable greater interaction with SIN3A, facilitating increased histone deacetylation to cope with elevated R-loop levels.

Based on these findings, we then wanted to directly examine THOC1’s role in regulating R-loop levels. Our data shows that THOC1-knockdown resulted in increased R-loop levels across multiple GBM lines, while THOC1 overexpression resulted in decreased R-loop levels in normal cells. Furthermore, THOC1-knockdown GBM cells also displayed decreased histone deacetylation activity, while THOC1-overexpression cells resulted in increased histone deacetylation activity – confirming the possibility that THOC1 may be regulating R-loop formation via modulation of the epigenetic landscape.

Given these results showing THOC1’s specific role in R-loop dynamics, we then wanted to explore whether certain networks were being selectively regulated by THOC1-mediated R-loop prevention. To explore this, we performed an unbiased RNA-sequencing screen to determine the identity of networks being uniquely governed by THOC1’s manipulation of R-loops. Intriguingly, we show that telomere packaging is specifically regulated by THOC1-mediated R-loop regulation, uncovering a novel dimension of THOC1’s involvement in GBM biology: telomere stability. THOC1-knockdown led to a striking increase in telomeric R-loop levels, which, in turn, was also found to be associated with significantly reduced telomere length. Conversely, THOC1-overexpression in normal cells resulted in decreased telomeric R-loop burden and significantly increased telomere lengths. These findings suggest that THOC1 may play a crucial role in preventing the formation of not only cellular R-loops in general, but specifically telomeric R-loops, known to be especially critical for the maintenance of functional telomeres.

Telomeric R-loops have recently gained attention for their potential impact on telomere stability and function. Several studies in the broader field of cancer biology have implicated telomeric R-loops in promoting genomic instability and cancer progression. Our data shows that increased telomeric R-loops as a result of depletion of THOC1 is associated with GBM cell death and shortened telomeres. Given that dysregulated telomeric R-loops have been shown to lead to telomere shortening and dysfunction in many other cancers, this suggests that THOC1 may be a promising target to artificially weaken telomeres and target GBM. Additionally, our data also shows a potential mechanism of THOC1-overexpression on transformation to a cancerous phenotype. Perhaps this is through decreased telomeric R-loops, which studies have previously shown to be involved in oncogenic transformation through telomerase inhibition. However, the mechanism of this still needs to be explored to understand this process fully.

Glioblastoma stands as one of the most aggressive and lethal cancers, characterized by an exceptionally poor prognosis despite advances in therapeutic approaches. We conducted a comprehensive CRISPR-Cas9 knockout screen, identifying approximately 150 previously unrecognized genes crucial for GBM progression. THOC1 was found to be significantly elevated in GBM and particularly responsible for promoting GBM aggressiveness. Our study illuminates the intricate landscape of GBM progression and oncogenesis by unveiling the critical role of THOC1. Previously, the precise workings of R-loops in GBM remained elusive; however, our findings illuminate how the delicate R-loop equilibrium is maintained by THOC1 in order to sustain replicative potential without leading to DNA damage. Here, we show the precise epigenetic mechanisms governing THOC1-mediated R-loop prevention and the downstream implications of this mechanism, including telomere maintenance. Overall, this study highlights a promising avenue for innovative GBM therapies and offers potential insights into combating cancer’s relentless progression. Specifically, targeting THOC1 emerges as an effective strategy to exploit R-loop levels and potentially induce GBM cell death, shedding light on the intricate connections between THOC1, R-loops, and GBM progression and paving the way for novel therapeutic strategies targeting the subtle yet fragile R-loop landscape.

## ACKNOWLEDGMENTS

This work was supported by the National Institute of Neurological Disorders and Stroke grant 1R01NS096376, 1R01NS112856, and P50CA221747 SPORE for Translational Approaches to Brain Cancer (to A.U.A.); 5R01NS110703, R01CA245969 and U19CA264338 (to A. M. S.). The results published here are partly based upon data generated by the TCGA Research Network: https://www.cancer.gov/tcga, and were further analyzed through GlioVis. In addition, these results use data generated by the Human Protein Atlas and GBMSeq (Gephart Lab, www.gbmseq.org). Figures, in part, were generated using BioRender (www.biorender.com).

---

**Figure S1:**
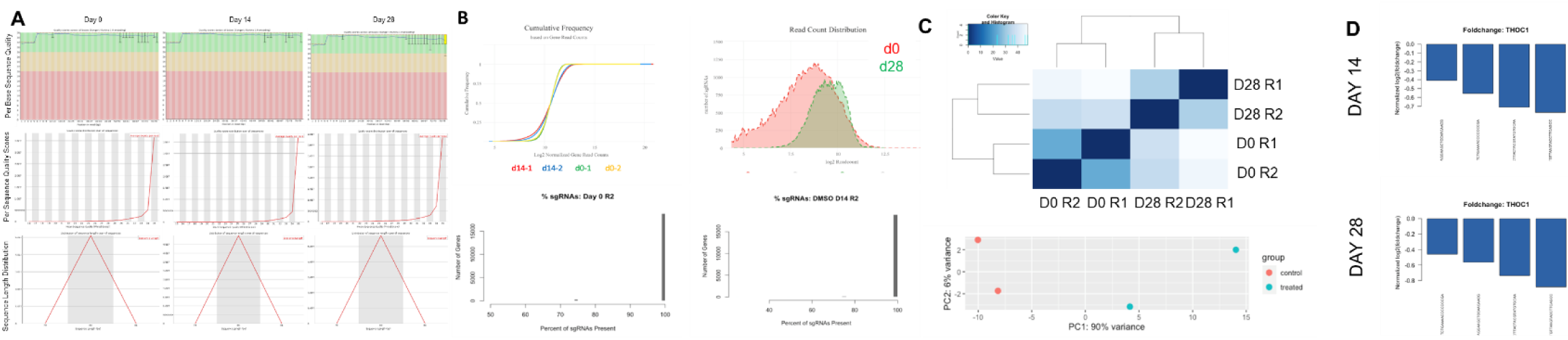
CRISPR-Cas9 screen quality. ***A)*** Quality assessment of sequencing data, displaying per base sequence quality, sequence quality scores, and length distribution to validate screen results. ***B)*** Distribution of gene read counts and sgRNA frequencies across conditions, confirming the screen’s reliability in identifying essential GBM drivers. ***C***) Principal component analysis (PCA) showing replicates’ consistency and condition differences, supporting the screen’s overall quality.

**Figure S2:**
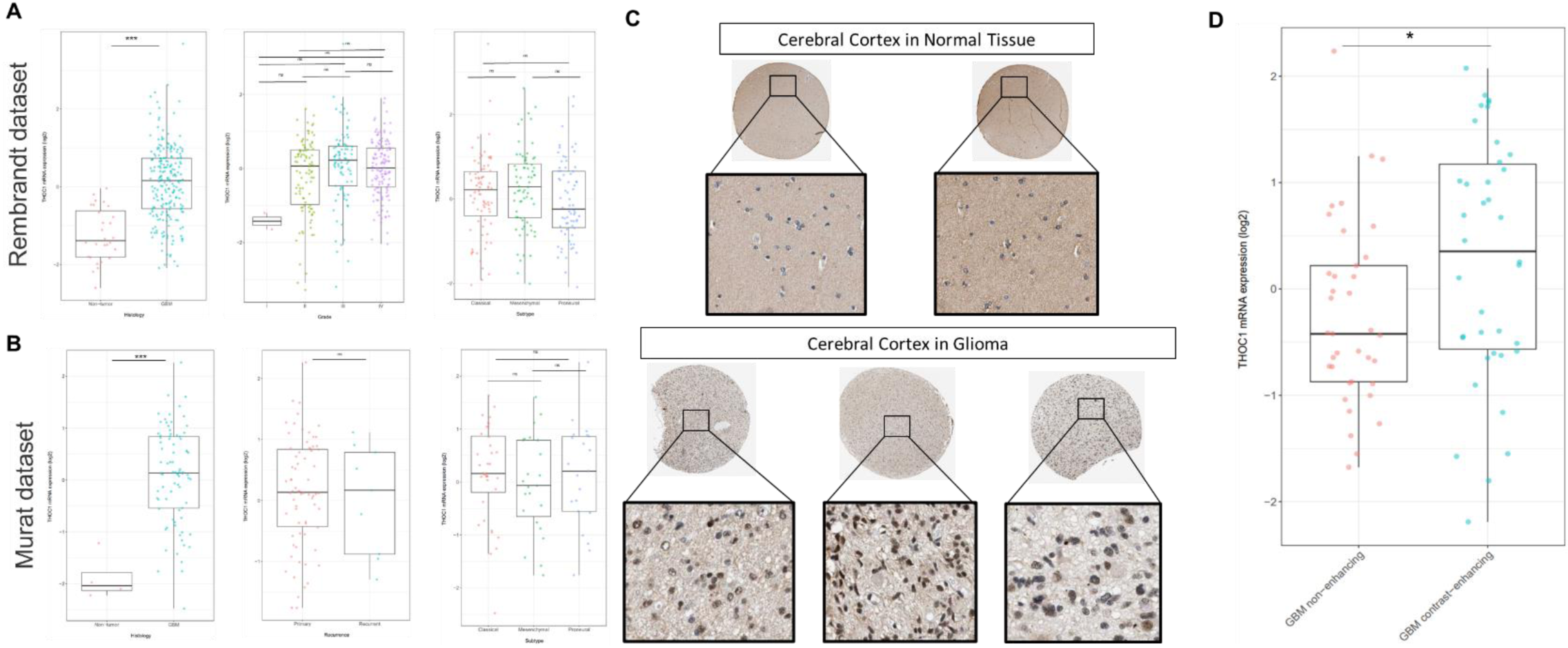
THOC1 expression in patient samples. ***A)*** Comparative analysis of THOC1 RNA expression across tumor grades, recurrence status, and subtypes showed no significant correlation, based on data from the GlioVis portal in the Rembrandt dataset. ***B)*** Comparative analysis of THOC1 RNA expression across tumor grades, recurrence status, and subtypes showed no significant correlation, based on data from the GlioVis portal in the Murat dataset. ***C)*** Qualitative assessment of THOC1 protein expression in GBM tissue compared to normal brain tissue from Protein Atlas. ***D)*** THOC1 RNA expression in non-enhancing versus contrast-enhancing regions of GBM from GlioVis portal.

**Figure S3:**
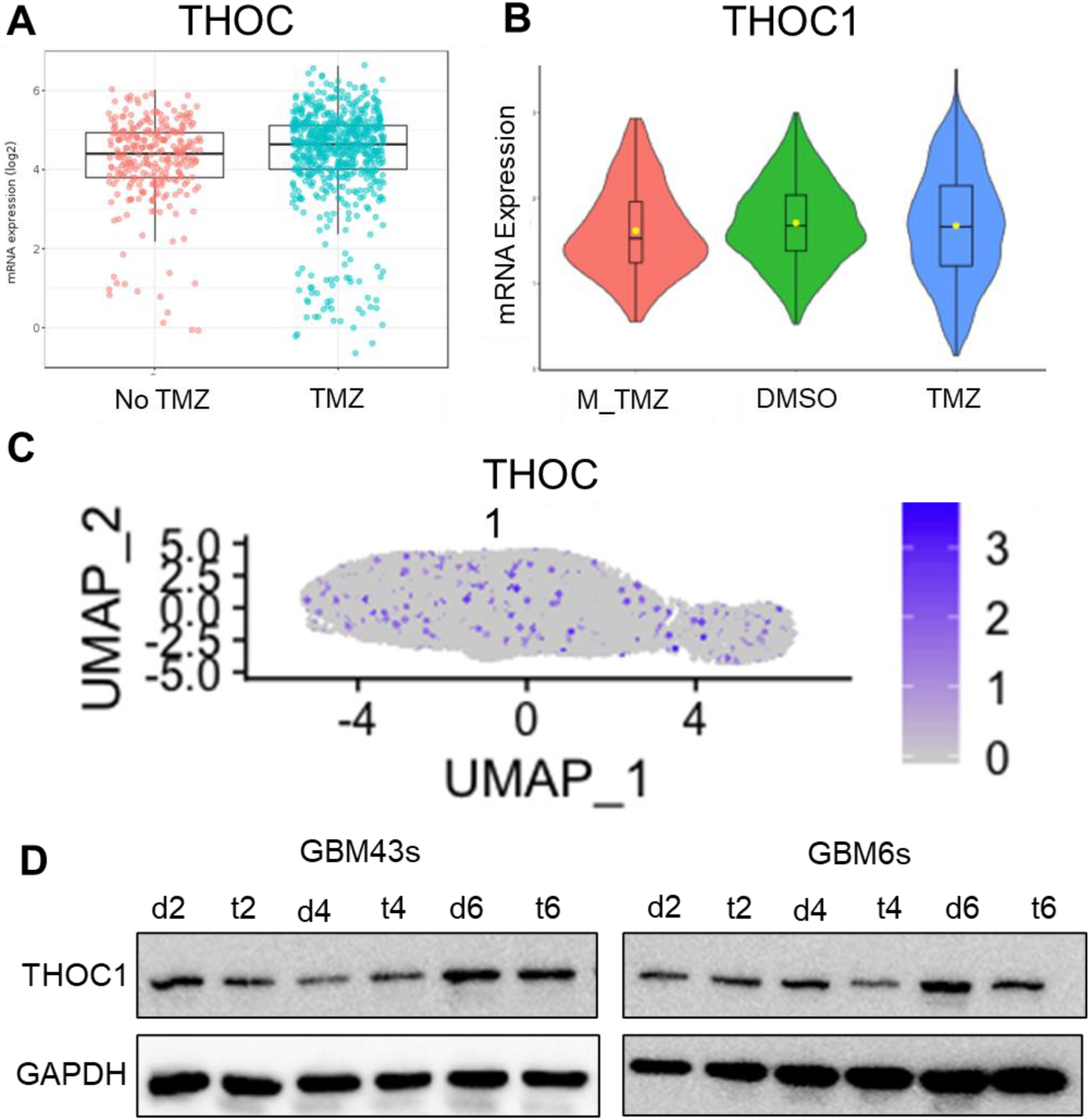
THOC1 Expression During TMZ Therapy in GBM. ***A)*** GlioVis portal analysis showing no increase in THOC1 RNA expression in GBM patients treated with temozolomide (TMZ) compared to those not treated. ***B) C)*** Single-cell RNA sequencing data comparing GBM cells post-DMSO, post-TMZ, and mid-therapy, demonstrating consistent THOC1 expression levels regardless of therapy status. ***D)*** Western blot analysis confirming no significant changes in THOC1 protein levels at various time points following TMZ exposure, indicating that THOC1 expression is stable during TMZ therapy.

**Figure S4:**
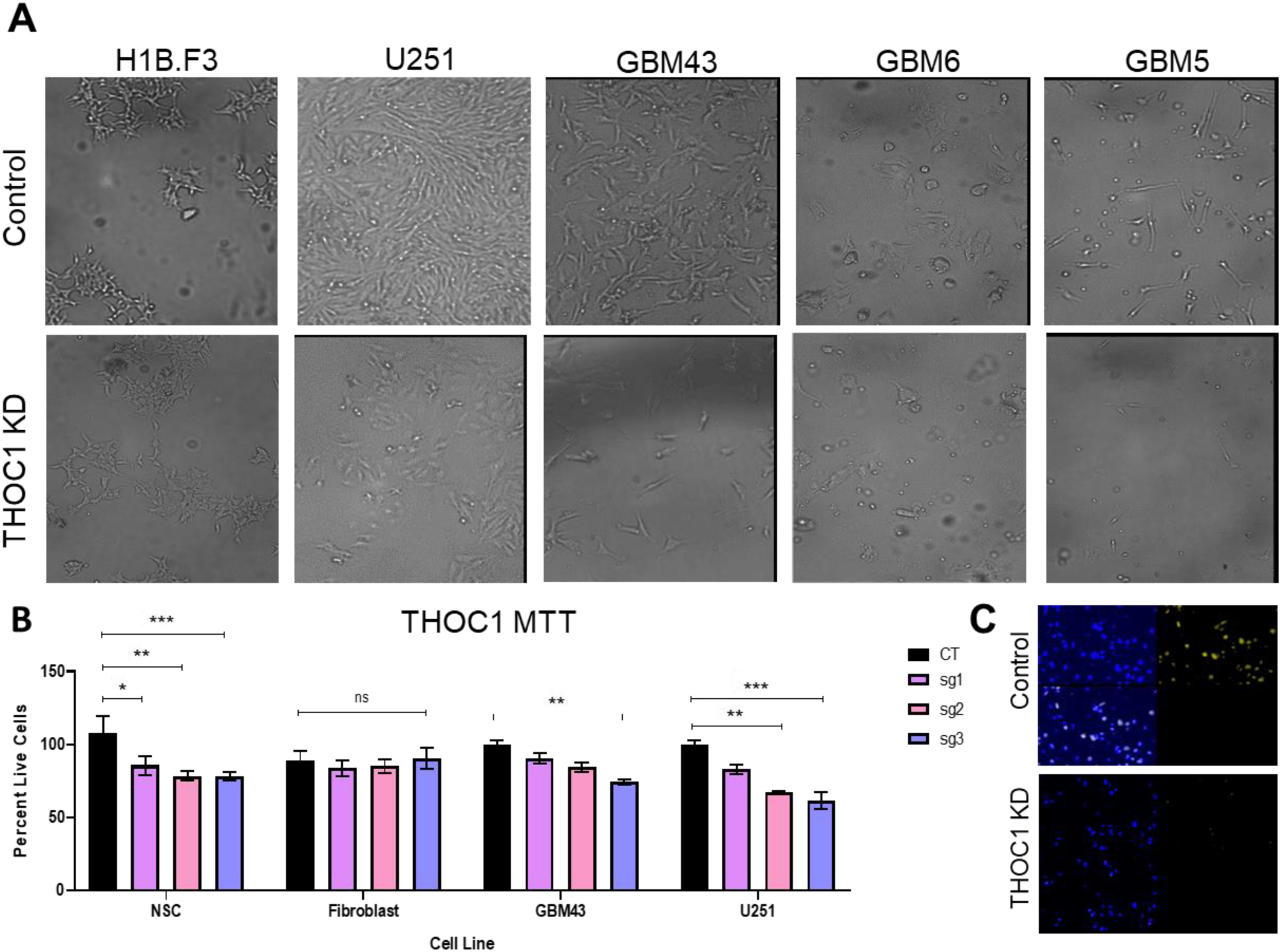
THOC1 Knockdown Reduces Viability and Proliferation in GBM Cells. ***A)*** Schematic showing the CRISPR-Cas9 guides used for THOC1 knockdown in selected GBM cell lines. ***B)*** Verification of decreased cell viability in THOC1 knockdown GBM cell lines through MTT assays. ***C)*** Additional MTT assay results showing reduced proliferation in THOC1-knockdown U251 and GBM43 cell lines, compared to knockdown fibroblasts. ***D)*** Immunohistochemical staining confirming THOC1 knockdown in brain tissue from mice implanted with THOC1-knockdown GBM43 cells.

**Figure S5:**
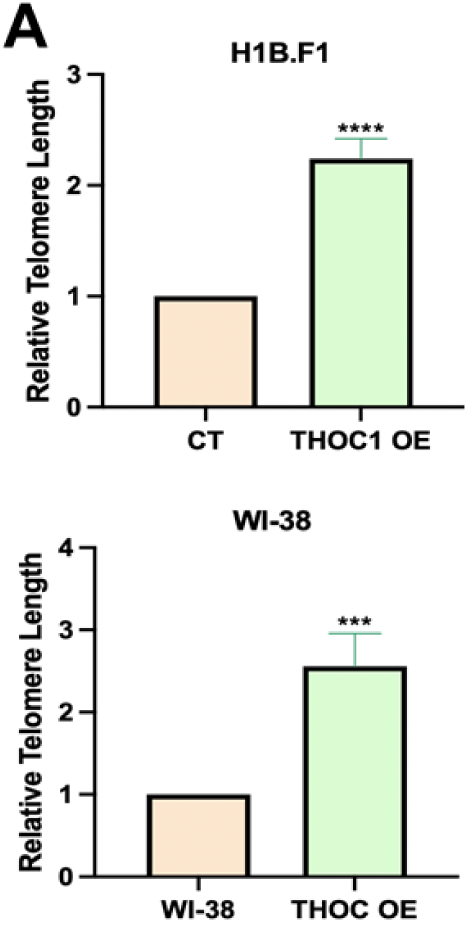
THOC1 Overexpression Results in Increased Telomere Length. ***A)*** THOC1 overexpression significantly increases telomere length in both H1B.F1 and WI-38 cells compared to controls.

